# Questioning the Evidence for Host-Symbiont Codiversification in Mycorrhizal Symbioses

**DOI:** 10.1101/2025.05.14.653600

**Authors:** Fantine Bodin, Hélène Morlon, Benoît Perez-Lamarque

**Affiliations:** Institut de Biologie de l’ENS (IBENS), École Normale Supérieure, CNRS, INSERM, Université PSL, 46 rue d’Ulm, 75005 Paris, France; Université de Toulouse, Toulouse INP, CNRS, IRD, CRBE, Toulouse, France

**Keywords:** Cophylogenetic signal, phylogenetic congruence, mycorrhizal symbioses, coevolution, codiversification, cospeciation

## Abstract

The vast majority of plants form mycorrhizal symbioses. The ecological importance of these mutualistic interactions has sparked interest in their long evolutionary history. By examining interaction networks and phylogenetic trees, several studies have suggested that plants codiversify with their associated mycorrhizal fungi. However, recent research has demonstrated that phylogenetic congruence (interpreted as codiversification when divergence times match) is often conflated with another pattern called cophylogenetic signal (*i.e.,* closely related plants interacting with closely related fungi and *vice versa*), which may arise from different biological processes. We performed cophylogenetic analyses on 29 diverse mycorrhizal networks to reevaluate the evidence for codiversification in mycorrhizal symbioses. We found significant cophylogenetic signal but no phylogenetic congruence: closely related plants often interact with closely related fungi, but their phylogenies do not match. Instead of codiversification, this finding is consistent with trait matching between plants and mycorrhizal fungi, where compatible, evolutionarily conserved plant and fungal traits govern their interactions. Our work highlights the importance of appropriately interpreting the cophylogenetic methods used to study the macroevolution of plants and their mycorrhizal fungi. It suggests that previous evidence of codiversification in mycorrhizal symbioses actually detected cophylogenetic signal, since across the multiple and diverse networks analyzed here, there is no evidence that codiversification occurred during the evolution of mycorrhizal symbioses.

## Main text

More than 90% of vascular plants form mycorrhizal symbioses (Brundrett and Tedersoo, 2018). These mutualistic associations are crucial for nutrition of most plants and their associated fungi and thus for the functioning of whole ecosystems (Smith and Read, 2008; Tedersoo et al., 2020; van der Heijden et al., 2015). Mycorrhizal symbioses originated more than 400 million years ago, likely facilitating the transition of plants from water to land (Remy et al., 1994; Strullu-Derrien et al., 2018). Over time, these associations evolved and diversified, leading to the recognition of several types of mycorrhizal symbioses (van der Heijden et al., 2015). The ancient origin and the ecological importance of these interactions have sparked interest in their assembly and evolution. In natural communities, mycorrhizal fungi typically interact with multiple plant species, rather than forming exclusive one-to-one associations (van der Heijden et al., 2015). Therefore, they shape complex interaction networks (Chagnon et al., 2012; Jacquemyn et al., 2015; Perez-Lamarque et al., 2022; Petrolli et al., 2022; Rimington et al., 2019; Sepp et al., 2019). These mycorrhizal networks are classical examples of bipartite ecological networks, representing interactions between species of two ecological guilds (Bascompte and Jordano, 2014). By considering the structure of mycorrhizal networks and the phylogenetic trees of plants and their associated mycorrhizal fungi, it has been suggested that phylogenetic congruence or even codiversification can occur in mycorrhizal symbioses (Arifin et al., 2023; Merckx and Bidartondo, 2008; Van Galen et al., 2023 ; see Table 1 for definitions). Whilst these studies discuss multiple processes by which these patterns could arise, one possibility is that plants and their associated mycorrhizal fungi could be undergoing simultaneous speciation events (*i.e.,* cospeciation; Table 2), leading to codiversification.

**Table 1:** Cophylogenetic definitions of evolutionary patterns in the context of mycorrhizal symbioses. These definitions follow the terminology of Perez-Lamarque and Morlon (2024). The patterns are ordered from the most general to the most specific.

| Pattern | Definition | Examples of generating processes |
| --- | --- | --- |
| **Cophylogenetic signal** | Tendency of closely related plant species to interact with closely related mycorrhizal fungal partners (and *vice versa*). | Trait matching;<br>Past biogeographic events |
| **Phylogenetic congruence** | Strong similarity between the phylogenetic trees of interacting plant and mycorrhizal fungal clades in terms of topology and relative branch lengths. | Diversification by preferential host-switching (De Vienne et al., 2013);<br>Phylogenetic tracking;<br>Successive vicariance events repeatedly affecting both clades |
| **Codiversification/ Co-cladogenesis** | Phylogenetic congruence between the plant and mycorrhizal fungal phylogenies, accompanied by matching divergence times. | Phylogenetic tracking;<br>Successive vicariance events repeatedly affecting both clades |

**Table 2:** Definitions of evolutionary processes in the context of mycorrhizal symbioses. These definitions follow the terminology of Perez-Lamarque and Morlon (2024).

| Process | Definition |
| --- | --- |
| **Coevolution** | Reciprocal evolutionary changes driven by selective pressures between two interacting plant and fungal lineages (pairwise coevolution). When more than two lineages are involved, the process is referred to as **diffuse coevolution**. Coevolution does not necessarily generate phylogenetic congruence. |
| **Cospeciation** | Concomitant speciation event in a plant lineage and its associated mycorrhizal fungal lineage. |
| **Phylogenetic tracking** | Repeated plant speciation events leading to subsequent speciations in their associated mycorrhizal fungal lineages (**cospeciations**), for example through vertical transmission that isolates fungal populations between diverging plant lineages. Fungal lineages may also undergo host transfers, intra-host duplications, or losses at low frequency. Phylogenetic tracking typically results in a pattern of codiversification. |
| **Trait matching** | Association of plants and fungi because of the complementarity of their mycorrhizal traits (for example plant root architecture, fungal nutrient exchange efficiency, or molecular signaling between plants and fungi). When these traits tend to be evolutionarily conserved, trait matching typically results in cophylogenetic signal. |
| **Vicariance** | Split of a population into two independently-evolving populations caused by the formation of a geographic barrier to gene flow (e.g. mountains, rivers, oceans). |

Testing for the occurrence of codiversification in host-symbiont associations is notoriously challenging, and has spurred the development of several cophylogenetic methods (Blasco-Costa et al., 2021; De Vienne et al., 2013; Dismukes et al., 2022; Page, 1994; Perez-Lamarque and Morlon, 2024; Ronquist, 1995). However, two distinct patterns – cophylogenetic signal and phylogenetic congruence – have often been conflated (Perez-Lamarque and Morlon, 2024) (Figure 1; Table 1). In the context of mycorrhizal symbioses, cophylogenetic signal indicates that closely related plants tend to interact with closely related mycorrhizal fungi and *vice versa*. In contrast, phylogenetic congruence is a more stringent, specific case of cophylogenetic signal, occurring when the phylogenetic trees of plants and their associated mycorrhizal fungi closely mirror each other (Figure 1). Codiversification refers to this pattern of phylogenetic congruence combined with matching divergence times, reflecting parallel diversification histories in plants and their associated fungi (De Vienne et al., 2013; Suzuki et al., 2022). Distinguishing these two patterns of cophylogenetic signal and phylogenetic congruence is crucial because they may stem from different biological processes. A cophylogenetic signal may simply arise if present-day mycorrhizal interactions assemble according to the trait matching of evolutionarily conserved traits (Perez-Lamarque and Morlon, 2024). It can also be generated by past biogeographic events that have constrained the distributions of extant species. In contrast, phylogenetic congruence arises from more stringent scenarios. For instance, codiversification can result from repeated cospeciation events driven by phylogenetic tracking, or from successive vicariance events that repeatedly affect both plants and their associated mycorrhizal fungi (Althoff et al., 2014; De Vienne et al., 2013; Perez-Lamarque and Morlon, 2024). These two patterns can be detected using two types of cophylogenetic methods, although they are not independent since phylogenetic congruence is a specific case of cophylogenetic signal. Global-fit methods such as ParaFit or PACo (Balbuena et al., 2013; Hutchinson et al., 2017; Legendre et al., 2002) can assess cophylogenetic signal, but only event-based methods such as Jane or eMPRess (Conow et al., 2010; Santichaivekin et al., 2021) can evaluate phylogenetic congruence.

**Figure 1:**
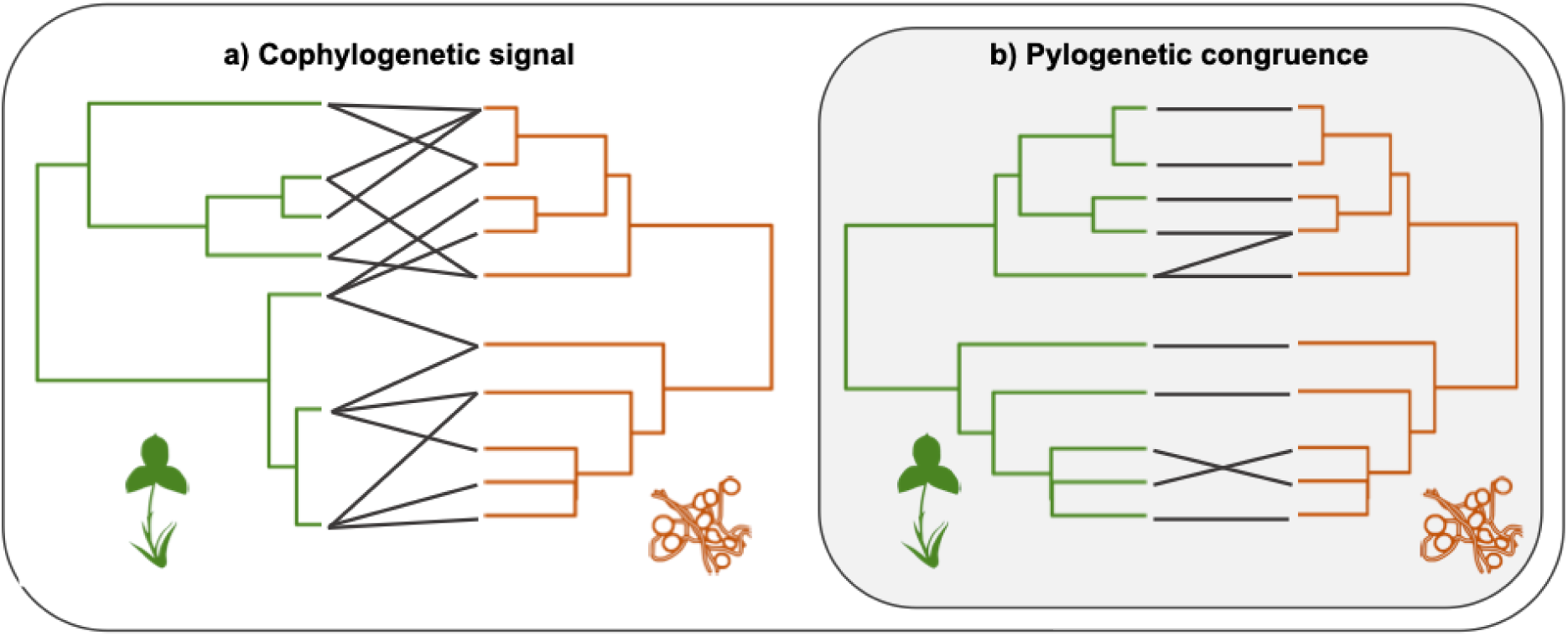
The patterns of cophylogenetic signal *versus* phylogenetic congruence. **a**. Cophylogenetic signal: closely related plants tend to interact with closely related mycorrhizal fungi and *vice versa*. **b.** Phylogenetic congruence: the phylogenetic trees of plants and their associated fungi mirror one another in terms of topology and relative branch lengths. When the divergence times of the two clades also coincide, this pattern corresponds to codiversification. The grey boxes indicate that phylogenetic congruence is a specific case of cophylogenetic signal (*i.e.,* phylogenetic congruence implies cophylogenetic signal, but the reverse is not necessarily true). See Table 1 for a list of definitions of cophylogenetic patterns and Table 2 for examples of generating processes.

Here, we reevaluate the evidence for codiversification in mycorrhizal symbioses in light of recent cophylogenetic developments (Perez-Lamarque and Morlon, 2024), using an extensive collection of mycorrhizal networks. We manually screened the literature and selected 29 networks from 13 different publications, ensuring coverage of various spatial and evolutionary scales, as well as all four main types of mycorrhizal symbioses (arbuscular mycorrhizae, ectomycorrhizae, orchid and ericoid mycorrhizae). For each mycorrhizal network, if the phylogenetic trees were not already available, we retrieved representative DNA sequences for both plant and mycorrhizal fungi, using multiple genes when possible (Supplementary Table 1). Sequences were aligned using MAFFT (Katoh, 2002) and refined using trimAl (Capella-Gutiérrez et al., 2009). Phylogenetic trees, as robust as possible, were then reconstructed using IQ-TREE (Nguyen et al., 2015), constraining deep nodes by a robust backbone tree when necessary (Supplementary Table 1). To assess cophylogenetic signal, we applied the global-fit method PACo (Balbuena et al., 2013; Hutchinson et al., 2017) to each mycorrhizal network. We then evaluated phylogenetic congruence using the event-based model eMPRess (Santichaivekin et al., 2021). This tool reconciles phylogenetic trees by fitting a phylogenetic-tracking model that accounts for events of cospeciation, transfer, loss, and duplication. eMPRess returns the number of each event and assesses the significance of the reconciled scenario through permutation tests. We considered phylogenetic congruence to be supported when the reconciled scenario had a significantly lower reconciliation cost than expected under permutation (p-value < 0.05) and included more cospeciation events than transfer events (Dorrell et al., 2021; Groussin et al., 2017; Perez-Lamarque and Morlon, 2024). Codiversification was directly discarded when no phylogenetic congruence was detected and, when phylogenetic congruence is present, can be further evaluated by comparing divergence times.

We detected a significant cophylogenetic signal in 13 out of 29 mycorrhizal networks (Figure 2). This pattern was consistently observed across the different types of mycorrhizal symbioses, regardless of the spatial and evolutionary scales (Supplementary Table 1). Yet, cophylogenetic signal was more frequently detected in some types of mycorrhizal symbioses (Fisher’s exact test: p=0.03, Supplementary Table 2): it was particularly significant in arbuscular mycorrhizal networks including mycoheterotrophic species (*i.e.* non-autotrophic, parasitic plants), and in ectomycorrhizal networks (Figure 2). However, more networks would be needed to test this pattern with sufficient statistical power. This result confirms that there is evidence of existing cophylogenetic signal in mycorrhizal associations. In contrast, phylogenetic congruence was absent in all 29 networks (Figure 2). This absence of phylogenetic congruence implies an absence of codiversification. Our results were consistent across phylogenetic reconstruction methods (Supplementary Table 1), meaning that uncertainty in fungal phylogenies does not explain the observed patterns. The fungal phylogenies of 4 of the 29 networks were built with only the ITS region, which may poorly resolve deep evolutionary relationships, but generally supports recent splits. Our results were robust despite uncertain deep phylogenetic resolution in some networks because recent splits play a major role in the detection of phylogenetic congruence, while incongruence at deeper nodes can often be reconciled by a few host-transfer events using eMPRess. Therefore, given the high diversity of mycorrhizal networks analyzed, the consistency of our results indicates that they might be generalized beyond the specific sets of species and systems included in our analyses. Our findings suggest that previous studies that proposed codiversification in mycorrhizal symbioses based on visual inspections or global-fit methods detected cophylogenetic signal rather than phylogenetic congruence, and that until now there is in fact no evidence of codiversification between plants and their associated mycorrhizal fungi in the networks analyzed here.

**Figure 2:**
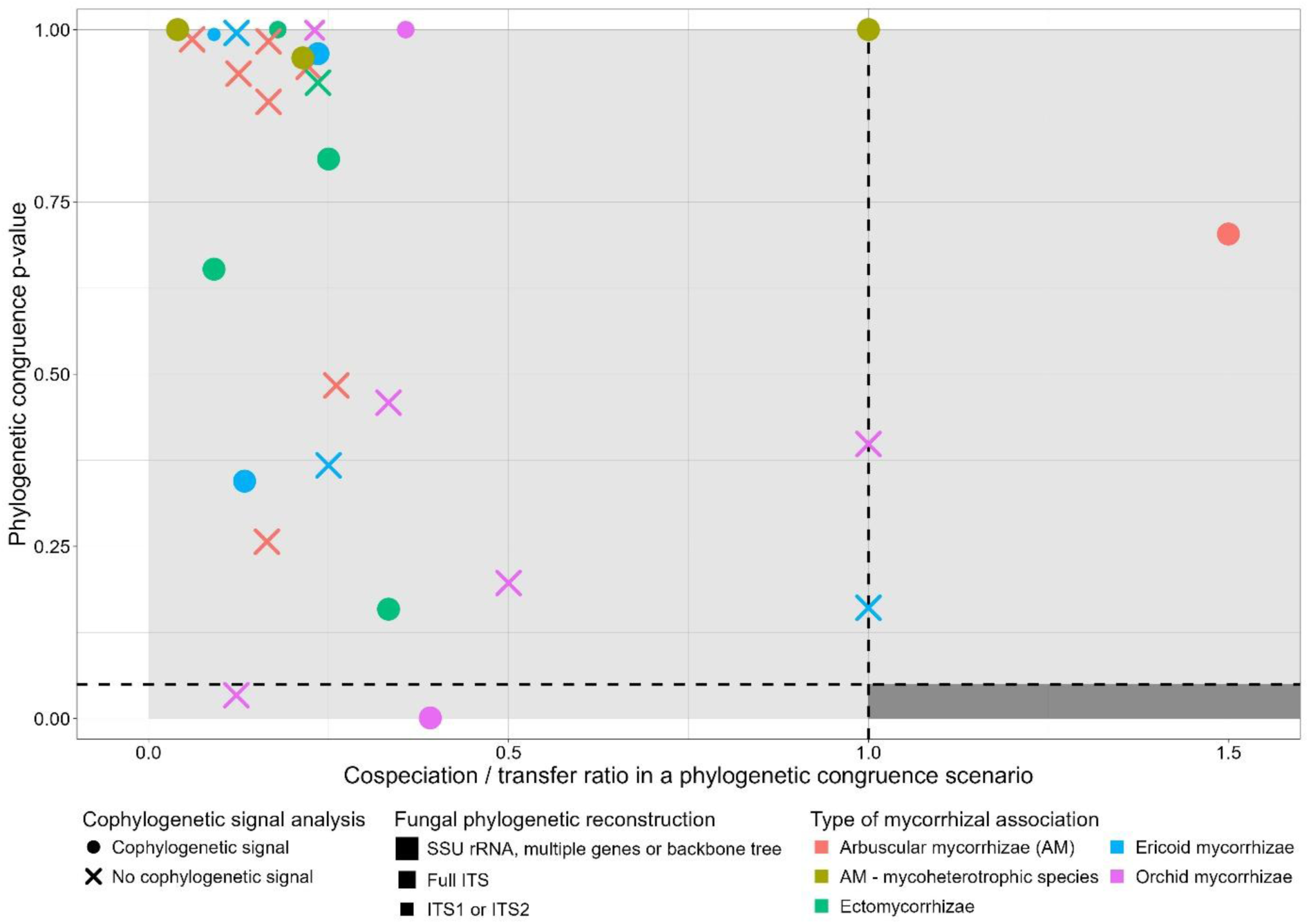
Cophylogenetic analysis of 29 mycorrhizal networks. For each mycorrhizal network, we report its eMPRess p-value as a function of the ratio of cospeciation to transfer events inferred by eMPRess. The different shades of grey indicate whether there is phylogenetic congruence: the dark grey area indicates significant phylogenetic congruence (eMPRess p-value < 0.05 and cospeciation-to-transfer ratio > 1), whereas the light grey areas indicate no phylogenetic congruence (eMPRess p-value > 0.05 and/or cospeciation-to-transfer ratio < 1). In addition, the shape of each symbol reflects the significance of the cophylogenetic signal inferred using PACo: a dot represents a network with significant cophylogenetic signal (PACo p-value < 0.05), whereas a cross represents a network without cophylogenetic signal (PACo p-value > 0.05). The size of each shape indicates the type of genetic data available for reconstructing the fungal phylogenetic trees and the color indicates the type of mycorrhizal symbiosis. Detailed information on each network can be obtained in Supplementary Table 1.

The absence of observed codiversification in mycorrhizal symbioses may stem from the fact that most mycorrhizal partners do not engage in highly specialized or one-to-one associations, but instead participate in complex interaction networks characterized by high partner sharing (Arifin et al., 2023; Hayward and Horton, 2014; Öpik et al., 2009; Perez-Lamarque et al., 2022; Sepp et al., 2019; Toju et al., 2016; Van Galen et al., 2023; Zhao et al., 2021). Such generalized interactions reduce the likelihood of frequent cospeciation (Althoff et al., 2014; Poisot, 2015), although our results do not rule out the occurrence of rare cospeciation events that may sporadically happen in some mycorrhizal associations and whose signal may be blurred by other events. While our findings suggest that codiversification is uncommon in mycorrhizal networks, this result also does not rule out the role of coevolution (Brundrett and Tedersoo, 2018; van der Heijden et al., 2015). The significant presence of cophylogenetic signal suggests that closely related plants tend to interact with closely related fungi. At regional and global scales, this signal could be at least partially explained by the biogeography of plants and/or mycorrhizal fungi. Vicariance events and dispersal limitations may lead to allopatric speciations and geographically independent radiations in both plants and fungi. As a result, closely related plants and closely related fungi tend to be present in similar places (Tedersoo et al., 2014), generating cophylogenetic signal (Perez-Lamarque and Morlon, 2024). However, we also observed cophylogenetic signal in mycorrhizal networks at local scale within individual communities (Supplementary Table 1), which cannot be due to biogeography. Therefore, the cophylogenetic signal we observed may reflect the fact that mycorrhizal interactions require a minimal level of trait matching, and that the involved mycorrhizal traits are evolutionarily conserved. This hypothesis is reinforced by the frequent detection of cophylogenetic signal in networks involving mycoheterotrophic plants and ectomycorrhizal associations, which are thought to require higher partner specificity driven by trait matching (Bruns, 2002; Merckx, 2012; Perez-Lamarque et al., 2020; Suetsugu et al., 2022; van der Heijden et al., 2015; Zhao et al., 2021). In this scenario, diffuse coevolution between interacting plants and fungi may have contributed to the evolutionary conservatism of key mycorrhizal traits (Cairney, 2000), such as plant root architecture, fungal nutrient exchange efficiency, or molecular signaling between plants and fungi (Castanedo et al., 2025; Delaux and Schornack, 2021; Karandashov et al., 2004; van der Heijden et al., 2015).

Our study highlights the importance of careful interpretation of cophylogenetic methods, and suggests that mycorrhizal symbionts did not codiversify with their host plants. Evolutionary history nevertheless plays an important role in shaping plant-mycorrhizal networks, as evidenced by the phylogenetic conservatism of interactions. Trait matching and trait conservatism may mediate this phylogenetic signal, suggesting that a deeper understanding of the functional traits underlying mycorrhizal compatibility is key to understanding the evolution of mycorrhizal symbioses. Finally, synthesizing our analyses with similar studies in other plant-associated interactions, such as pollination, seed dispersal, or parasitic interactions (Asar et al., 2022; Jackson, 2004; Russo et al., 2018; Taengon et al., 2025), would help clarify whether codiversification has ever played a significant role in the evolutionary history of land plants.

## Methods

### Screening for available mycorrhizal networks

We manually screened the literature to compile as many mycorrhizal networks as possible (Supplementary Table 1). We targeted publications that provided freely accessible interaction networks, along with the phylogenetic trees or the DNA sequences of both plants and mycorrhizal fungi. We retained networks from natural habitats and with taxonomic delineations at the plant species level. Fungal operational taxonomic units (OTUs) were typically delineated using a 97% similarity threshold. We aimed to include all major types of mycorrhizal symbioses: arbuscular mycorrhizae (n=11), including three arbuscular mycorrhizal networks with mycoheterotrophic plant species (*i.e.* non-photosynthetic plant species that rely on mycorrhizal fungi for both organic and mineral nutrition), ectomycorrhizae (n=5), orchid mycorrhizae (n=7), and ericoid mycorrhizae (n=6). We also selected networks encompassing a broad range of spatial scales – from communities in specific habitats of La Réunion (Perez-Lamarque et al., 2022) to the global arbuscular mycorrhizal network between Glomeromycotina fungi and land plant species (Öpik et al., 2010; Perez-Lamarque et al., 2020) – and evolutionary scales – from a 14 million-year-old ectomycorrhizal association between Pisonieae and their thelephoroid fungal symbionts (Hayward and Horton, 2014) to more than 450 million years of diversification of all land plants forming arbuscular mycorrhizae. Our networks also encompass a diversity of sampling strategies: from sampling all the species present in a local community (for example Perez-Lamarque et al., 2022; Sepp et al., 2019), to intentionally focusing only on some specific plant clades (as in Arifin et al., 2023; Shefferson et al., 2010) or gathering all plant-fungal associations available at regional or global scale (for instance Öpik et al., 2009; Perez-Lamarque et al., 2020). In total, we studied 29 networks from 13 different publications (Arifin et al., 2023; Hayward and Horton, 2014; Martos et al., 2012; Merckx and Bidartondo, 2008; Öpik et al., 2010, 2009; Otero et al., 2011; Perez-Lamarque et al., 2022; Perez-Lamarque et al., 2020; Sepp et al., 2019; Shefferson et al., 2010; Toju et al., 2016; Van Galen et al., 2023; Zhao et al., 2021).

### Recovering phylogenetic trees

For some publications, phylogenetic trees were already publicly available (Supplementary Table 1). For the others, we retrieved DNA sequences from GenBank, aligned them with MAFFT (Katoh, 2002), trimmed them with trimAl (Capella-Gutiérrez et al., 2009), and then reconstructed the phylogenetic trees with IQ-TREE (Nguyen et al., 2015). When multiple sequences were available for a single species, we selected one representative (Supplementary Table 3). For the plants, we consistently used sequences from at least two genes, selected from trnL, ITS, matK, matR, 18S, and ATPa (Supplementary Table 1). When no genetic data were available in the original publications, we reconstructed the missing plant phylogenetic trees with V.PhyloMaker2 (Jin and Qian, 2022). For the fungal phylogenetic reconstructions, the available sequences primarily included the SSU rRNA gene and/or the ITS region (ITS1, ITS2, or the full ITS1-5.8S-ITS2 fragment), sometimes in addition to other genes (28S, EF1 alpha, TOR2, ML 4-5). The phylogenies obtained with multiple genes were considered robust. While the SSU rRNA gene alone can support robust phylogenetic trees, the fungal ITS region is often too variable, especially ITS1 or ITS2 alone, to reliably resolve deeper evolutionary relationships (Perez-Lamarque et al., 2022; Schoch et al., 2012). Therefore, to obtain as robust as possible phylogenetic trees when only sequences of partial ITS1 or ITS2 were available, we informed the IQ-TREE reconstructions with a robust backbone tree built from multiple-gene datasets (following Talavera et al., 2022). Specifically, we used the backbone trees of Weiß et al. (2016) for Sebacinales, of Johnston et al. (2019) for Helotiales, and of Zhang et al. (2025) for Cantharellales (Supplementary Table 1). This backbone approach improves bootstrap branch support, raising values >70% for most nodes. In total, 15 fungal phylogenetic trees were reconstructed using either multi-gene datasets or a backbone-tree approach, 10 using the SSU rRNA gene, 3 using the full ITS region, and 1 using only ITS1, as no additional sequences or backbone trees were available for this mycorrhizal system (Supplementary Table 1).

### Evaluating cophylogenetic signal

We assessed cophylogenetic signal, *i.e.* whether closely related plants tend to interact with closely related fungi, using the global-fit method PACo (Balbuena et al., 2013; Hutchinson et al., 2017). This method returns a R^2^ and a p-value obtained by comparison with permutations. We ran 10,000 permutations for each mycorrhizal network. Additionally, we checked the results with the global-fit method ParaFit (Legendre et al., 2002; Paradis et al., 2004). Both methods yielded similar results (Supplementary Table 1). We then tested whether the type of mycorrhizal symbioses influences the significance of the cophylogenetic signal using a Fisher’s exact test.

### Evaluating phylogenetic congruence

We assessed phylogenetic congruence, *i.e.* whether the phylogenetic trees of plants and mycorrhizal fungi tend to mirror one another, with the event-based model eMPRess (Santichaivekin et al., 2021). eMPRess assumes a process of phylogenetic tracking and tries to reconcile the plant and fungal phylogenetic trees by fitting, using maximum parsimony, events of cospeciation, host transfer, duplication, and loss, with a given cost associated to each event. We ran eMPRess with the following costs: 0 for a cospeciation event, 4 for a duplication event, 1 for a loss event, and 1 for a transfer event. We obtained similar results using different cost values (Supplementary Table 1). To assess the significance of each eMPRess reconciliation, we performed 1,000 permutations of the mycorrhizal associations. We considered a reconciliation as significant when the total cost was lower than 95% of the costs obtained from the permuted data (eMPRess p-value < 0.05) and when the number of cospeciation events was larger than the number of transfer events (cospeciation-to-transfer ratio > 1 (Perez-Lamarque and Morlon, 2024)). Indeed, transfer events can obscure cospeciation signals and generate a symbiont phylogeny that no longer matches the host phylogeny.

Importantly, eMPRess, like most event-based methods, evaluates only phylogenetic congruence because they consider only the topologies of the plant and fungal phylogenetic trees. It does not take into account whether the divergence times of corresponding plant and fungal nodes are concomitant to fit a cospeciation event. Thus, to conclude that codiversification has occurred, both a significant phylogenetic congruence (as detected by eMPRess) and coincident divergence times between the two clades are needed (De Vienne et al., 2013). In this study, because we did not detect significant phylogenetic congruence in any network, we did not need to evaluate coincidence of divergence times between the two clades to rule out codiversification.

As phylogenetic congruence can only exist if one-to-one host-symbiont interactions are frequent, eMPRess cannot account for symbionts interacting with multiple hosts, like most event-based approaches (De Vienne et al., 2013; Dismukes et al., 2022). Therefore, when a fungal species interacted with several plant species, we considered two approaches (Perez-Lamarque and Morlon, 2024) : (1) we only picked the most abundant interaction (or one interaction at random when abundances were not available; Supplementary Table 1), or (2) we randomly simulated bifurcating sub-trees for fungal taxa with multiple plants, such that each tip in the fungal tree is associated with a single plant tip (Satler et al., 2019). As the second approach did not change our findings, we only reported the results of the first approach in the main text (Supplementary Table 1).

It has been demonstrated using simulations that eMPRess still performs well when the number of codiversifying species is low, with statistical power exceeding 85% when the number of species is between 10 and 50 (Perez-Lamarque and Morlon, 2024). Thus, our results are robust to the small sample sizes of some networks (average number of species per clade = 14; Supplementary Table 1).

Finally, we verified that our results were robust to the phylogenetic uncertainty in the fungal tree by running eMPRess with different fungal phylogenetic trees obtained with different reconstruction methods (ITS only or the backbone approach). We obtained consistent results in all cases.

## Acknowledgments

The authors acknowledge members of the ‘Modeling biodiversity’ lab at IBENS and Laura Van Galen for helpful discussions, as well as Alice Garcia for initiating the project with preliminary analyses.

The authors also acknowledge Arild Arifin and Zhongtao Zhao for sharing the interaction networks and phylogenetic trees of plants and mycorrhizal fungi they studied, as well as all other authors whose data were directly available in their publications.

## Author contributions

FB, HM, and BPL designed the study, FB performed the analyses, FB and BPL wrote a first version of the manuscript and all authors contributed to the revisions.

## Code availability

All the scripts for performing the analyses, the phylogenetic tree files of fungi and plants, the DNA alignments used to reconstruct the trees, and the mycorrhizal networks are available through the following link: https://github.com/fantine-bodin/Cophylogenetics_mycorrhizal-symbioses.git.

## Supporting information

**Supplementary Table 1: Summary table with characteristics of each mycorrhizal network and the corresponding results of the cophylogenetic analyses.** The table reports, for each network: the spatial, evolutionary, and taxonomic scales, the network size, the type of mycorrhizal associations, how we recovered phylogenetic trees of both plants and fungi, the results of the evaluation of cophylogenetic signal (with 2 methods: PACo and ParaFit), and the results of the evaluation of phylogenetic congruence (with the event-based method eMPRess with different cost values for each reconciliation event).

**Supplementary Table 2: Results of cophylogenetic signal depending on the type of mycorrhizal symbioses.**

**Supplementary Table 3: Data selection for the mycorrhizal networks from Arifin et al. (Arifin et al., 2023).** Modification of Table S1 from Arifin et al. (2023). As different sequences of the same OTU were available, the representative sequence of each fungal OTU chosen to build the phylogenetic tree are highlighted in green.

