## Supplementary material for "Questioning the Evidence for Host-Symbiont Codiversification in Mycorrhizal Symbioses": Supp Table 2

**Supplementary Table 2: Results of cophylogenetic signal depending on the type of mycorrhizal symbioses.**

Cophylogenetic signal was assessed with PACo (significant signal if p-value < 0.05).

| **Mycorrhizal type** | **Significant cophylogenetic signal** | **Non-significant cophylogenetic signal** | **Total** |
|---|---|---|---|
| Arbuscular mycorrhizae (AM) | 1 | 7 | 8 |
| AM including mycoheterotrophic plant species | 3 | 0 | 3 |
| Ectomycorrhizae | 4 | 1 | 5 |
| Ericoid mycorrhizae | 3 | 3 | 6 |
| Orchid mycorrhizae | 2 | 5 | 7 |
| **Total** | 13 | 16 | 29 |
