## Supplementary material for "Questioning the Evidence for Host-Symbiont Codiversification in Mycorrhizal Symbioses": Supp Table 3

Supplementary Table 2. Modification of Table S1 from *Arifin, A.R., Phillips, R.D., Linde, C.C., 2023. Strong phylogenetic congruence between Tulasnella fungi and their associated Drakaeinae orchids. J. Evol. Biol. 36, 221–237. https://doi.org/10.1111/jeb.14107*. As different sequences of the same OTU were available, the representative sequence of each fungal OTU chosen to build the phylogenetic tree are highlighted in green.

| *Tulasnella* species/OTU | GenBank Number | Orchid host / substrates | Isolate / clone code | Study |
|---|---|---|---|---|
| *Tulasnella densa* | JN015192 | *Cryptostylis hunteriana* | BB0002_2_A | Howard & Clements 2011* |
| *Tulasnella densa* | MT036526 | *Cryptostylis hunteriana* | CLM2110 | Arifin et al. 2021 |
| *Tulasnella densa* | MT036524 | *Cryptostylis hunteriana* | CLM2112 | Arifin et al. 2021 |
| *Tulasnella densa* | MT036519 | *Cryptostylis hunteriana* | CLM2118 | Arifin et al. 2021 |
| *Tulasnella densa* | MT327885 | *Cryptostylis erecta* | CL145_5_clone1 | Unpublished |
| OTU 1 | OM311932 | *Arthrochilus corinnae* | CL153.2 clone1 | This study |
| OTU 1 | OM311933 | *Arthrochilus corinnae* | CL153.2 clone2 | This study |
| OTU 1 | OM311934 | *Arthrochilus corinnae* | CL153.2 clone3 | This study |
| OTU 1 | OM311935 | *Arthrochilus corinnae* | CL153.4 clone1 | This study |
| OTU 1 | OM311936 | *Arthrochilus corinnae* | CL153.4 clone2 | This study |
| OTU 1 | ON256304 | *Arthrochilus corinnae* | CLM2245 | This study |
| OTU 1 | ON256305 | *Arthrochilus corinnae* | CLM2246 | This study |
| OTU 1 | ON256306 | *Arthrochilus corinnae* | CLM2247 | This study |
| OTU 1 | ON256307 | *Arthrochilus corinnae* | CLM2248 | This study |
| *Tulasnella concentrica* | MT036547 | *Cryptostylis leptochila* | CLM2071 | Arifin et al. 2021 |
| *Tulasnella concentrica* | MT036546 | *Cryptostylis leptochila* | CLM2072 | Arifin et al. 2021 |
| *Tulasnella concentrica* | MT036545 | *Cryptostylis leptochila* | CLM2142 | Arifin et al. 2021 |
| *Tulasnella concentrica* | MT036532 | *Cryptostylis leptochila* | CLM2199 | Arifin et al. 2021 |
| *Tulasnella prima* | HM196783 | *Chiloglottis* aff. *jeanesii* | CP0835.VIII.2 | Roche et al. 2010 |
| *Tulasnella prima* | HM196784 | *Chiloglottis* aff. *jeanesii* | CP0835.VIII.2 | Roche et al. 2010 |
| *Tulasnella prima* | HM196788 | *Chiloglottis* aff. *jeanesii* | CP0835.III.1 | Roche et al. 2010 |
| *Tulasnella prima* | HM196786 | *Chiloglottis* aff. *jeanesii* | CP0835.IX.2 | Roche et al. 2010 |
| *Tulasnella prima* | HM196779 | *Chiloglottis* aff. *jeanesii* | CP0835.III.1 | Roche et al. 2010 |
| *Tulasnella prima* | HM196785 | *Chiloglottis* aff. *jeanesii* | CP0835.IX.2 | Roche et al. 2010 |
| *Tulasnella prima* | HM196782 | *Chiloglottis* aff. *jeanesii* | CP0835.I.1 | Roche et al. 2010 |
| *Tulasnella prima* | HM196795 | *Chiloglottis* aff. *jeanesii* | CP0835.IX.1 | Roche et al. 2010 |
| *Tulasnella prima* | HM196787 | *Chiloglottis* aff. *jeanesii* | CP0835.VIII.2 | Roche et al. 2010 |
| *Tulasnella prima* | HM196791 | *Chiloglottis* aff. *jeanesii* | 06082.I.1 | Roche et al. 2010 |
| *Tulasnella prima* | HM196792 | *Chiloglottis* aff. *jeanesii* | 06082.I.1 | Roche et al. 2010 |
| *Tulasnella prima* | KF476543 | *Chiloglottis formicifera* | CLM309 | Roche et al. 2010 |
| *Tulasnella prima* | KF476550 | *Chiloglottis formicifera* | CLM306 | Roche et al. 2010 |

| | | | | |
|---|---|---|---|---|
| *Tulasnella prima* | KF476551 | *Chiloglottis formicifera* | CLM308 | Roche et al. 2010 |
| *Tulasnella prima* | HM196796 | *Chiloglottis trapeziformis* | SRBG01.II.1 | Roche et al. 2010 |
| *Tulasnella prima* | HM196810 | *Chiloglottis trapeziformis* | SRBG03.III.12 | Roche et al. 2010 |
| *Tulasnella prima* | HM196799 | *Chiloglottis trapeziformis* | CM07.I.10 | Roche et al. 2010 |
| *Tulasnella prima* | HM196794 | *Chiloglottis trapeziformis* | CM07.I.5 | Roche et al. 2010 |
| *Tulasnella prima* | HM196806 | *Chiloglottis trapeziformis* | SRBG03.IV.5 | Roche et al. 2010 |
| *Tulasnella prima* | HM196809 | *Chiloglottis trapeziformis* | SRBM01.I.3.1 | Roche et al. 2010 |
| *Tulasnella prima* | HM196793 | *Chiloglottis trapeziformis* | SRBG01.II.3 | Roche et al. 2010 |
| *Tulasnella prima* | HM196807 | *Chiloglottis trapeziformis* | SRBG03.I.8 | Roche et al. 2010 |
| *Tulasnella prima* | HM196790 | *Chiloglottis trapeziformis* | SRBG01.II.3 | Roche et al. 2010 |
| *Tulasnella prima* | HM196789 | *Chiloglottis trapeziformis* | CM07.II.1 | Roche et al. 2010 |
| *Tulasnella prima* | HM196797 | *Chiloglottis seminuda* | 07033.II.2 | Roche et al. 2010 |
| *Tulasnella prima* | HM196798 | *Chiloglottis seminuda* | 07033.II.2 | Roche et al. 2010 |
| *Tulasnella prima* | HM196800 | *Chiloglottis seminuda* | 07033-45.II.2 | Roche et al. 2010 |
| *Tulasnella prima* | MK430473 | *Cryptostylis ovata* | CL13(6) | Nguyen et al. 2019 |
| *Tulasnella prima* | MK430470 | *Cryptostylis ovata* | CL12(5) | Nguyen et al. 2019 |
| *Tulasnella prima* | MK430521 | *Cryptostylis ovata* | CS43(17) | Nguyen et al. 2019 |
| *Tulasnella prima* | HM196801 | *Chiloglottis valida* | CV0836.I.1 | Roche et al. 2010 |
| *Tulasnella prima* | HM196804 | *Chiloglottis valida* | CV0627.II.1 | Roche et al. 2010 |
| *Tulasnella prima* | KF476556 | *Chiloglottis trilabra* | CLM159 | Roche et al. 2010 |
| *Tulasnella prima* | HM196805 | *Chiloglottis reflexa* | 07061.I.1 | Roche et al. 2010 |
| *Tulasnella prima* | HM196803 | *Chiloglottis diphylla* | 505.III.5 | Roche et al. 2010 |
| *Tulasnella prima* | KF476552 | *Chiloglottis diphylla* | CLM068 | Roche et al. 2010 |
| *Tulasnella prima* | MZ576545 | *Paracaleana nigrita* | CL103.3 clone1 | This study |
| *Tulasnella prima* | MZ576546 | *Paracaleana nigrita* | CL103.3 clone2 | This study |
| *Tulasnella prima* | MZ576547 | *Paracaleana nigrita* | CL103.3 clone3 | This study |
| *Tulasnella prima* | MZ576548 | *Paracaleana nigrita* | CL103.3 clone4 | This study |
| *Tulasnella prima* | MZ576549 | *Paracaleana nigrita* | CL103.3 clone5 | This study |
| *Tulasnella occidentalis* | MK430439 | *Cryptostylis ovata* | CB14(1) | Nguyen et al. 2020 |
| *Tulasnella occidentalis* | MK430440 | *Cryptostylis ovata* | CB14(3) | Nguyen et al. 2020 |
| *Tulasnella occidentalis* | MK430444 | *Cryptostylis ovata* | CB43(6) | Nguyen et al. 2020 |
| *Tulasnella occidentalis* | MK430445 | *Cryptostylis ovata* | CB43(9) | Nguyen et al. 2020 |
| *Tulasnella occidentalis* | MT008093 | *Cryptostylis ovata* | CLM1941 | Arifin et al. 2021 |
| *Tulasnella occidentalis* | MT008092 | *Cryptostylis ovata* | CLM1942 | Arifin et al. 2021 |
| *Tulasnella occidentalis* | MT008091 | *Cryptostylis ovata* | CLM1943 | Arifin et al. 2021 |
| *Tulasnella* sp. Jarrahdale 1 | MK430451 | *Cryptostylis ovata* | CJ11(8) | Nguyen et al. 2020 |
| *Tulasnella* sp. Jarrahdale 1 | MK430452 | *Cryptostylis ovata* | CJ13(3) | Nguyen et al. 2020 |
| *Tulasnella* sp. Jarrahdale 2 | MK430461 | *Cryptostylis ovata* | CJ51(7) | Nguyen et al. 2020 |
| *Tulasnella* sp. Jarrahdale 2 | MK430468 | *Cryptostylis ovata* | CJ51(17) | Nguyen et al. 2020 |
| *Tulasnella punctata* | MT008119 | *Cryptostylis subulata* | CLM2020 | Arifin et al. 2021 |
| *Tulasnella punctata* | MT008122 | *Cryptostylis subulata* | CLM2017 | Arifin et al. 2021 |
| *Tulasnella punctata* | MT008120 | *Cryptostylis subulata* | CLM2019 | Arifin et al. 2021 |

| | | | | |
|---|---|---|---|---|
| *Tulasnella punctata* | MT008108 | *Cryptostylis subulata* | CLM2031 | Arifin et al. 2021 |
| *Tulasnella punctata* | MT008107 | *Cryptostylis subulata* | CLM2032 | Arifin et al. 2021 |
| *Tulsnella sphagneti* | MT214495 | *Cryptostylis subulata* | CLM2131 | Arifin et al. 2021 |
| *Tulsnella sphagneti* | MT214496 | *Cryptostylis subulata* | CLM2132 | Arifin et al. 2021 |
| *Tulsnella sphagneti* | MT214493 | *Cryptostylis subulata* | CLM1952 | Arifin et al. 2021 |
| *Tulsnella sphagneti* | KY445922 | *Chiloglottis turfosa* | 12030 | Roche et al. 2010 |
| *Tulsnella sphagneti* | KY445924 | *Chiloglottis turfosa* | 13102.1 | Roche et al. 2010 |
| *Tulsnella sphagneti* | KY445923 | *Chiloglottis turfosa* | 13102.2 | Roche et al. 2010 |
| *Tulsnella sphagneti* | KY445926 | *Chiloglottis* sp. | 13065.1 | Roche et al. 2010 |
| *Tulsnella sphagneti* | KY445925 | *Chiloglottis* sp. | 13065.2 | Roche et al. 2010 |
| *Tulsnella sphagneti* | KY095117 | *Chiloglottis* aff. *valida* | 12033.1 | Roche et al. 2010 |
| *Tulsnella sphagneti* | KY445927 | *Chiloglottis* sp. | 13058 | Roche et al. 2010 |
| *Tulsnella sphagneti* | KY445928 | *Chiloglottis* sp. | 13139 1 | Roche et al. 2010 |
| *Tulsnella sphagneti* | KY445929 | *Chiloglottis* sp. | 13143 1 | Roche et al. 2010 |
| *Tulasnella eichleriana* | KC152389 | Decayed wood | DC294 | Cruz et al. 2014 |
| *Tulasnella eichleriana* | AY373292 | Unknown | KC852 | Cruz et al. 2014 |
| *Tulasnella rosea* | JX138568 | *Spiculaea ciliata* | MB-2012 | Martos et al. 2012 |
| *Tulasnella rosea* | MN947568 | *Spiculaea ciliata* | CLM1770 | Arifin et al. 2021 |
| *Tulasnella rosea* | MN947569 | *Spiculaea ciliata* | CLM1773 | Arifin et al. 2021 |
| *Tulasnella rosea* | MN947570 | *Spiculaea ciliata* | CLM1774 | Arifin et al. 2021 |
| *Tulasnella rosea* | MN947571 | *Spiculaea ciliata* | CLM1775 | Arifin et al. 2021 |
| *Tulasnella rosea* | MN947565 | *Spiculaea ciliata* | CLM1763 | Arifin et al. 2021 |
| *Tulasnella rosea* | MN947566 | *Spiculaea ciliata* | CLM1765 | Arifin et al. 2021 |
| *Tulasnella rosea* | MN947563 | *Spiculaea ciliata* | CLM1791 | Arifin et al. 2021 |
| *Tulasnella rosea* | MN947562 | *Spiculaea ciliata* | CLM1792 | Arifin et al. 2021 |
| *Tulasnella rosea* | MN947572 | *Spiculaea ciliata* | CLM1771 | Arifin et al. 2021 |
| *Tulasnella rosea* | MN947573 | *Spiculaea ciliata* | CLM1776 | Arifin et al. 2021 |
| *Tulasnella rosea* | MN947561 | *Spiculaea ciliata* | CLM1796 | Arifin et al. 2021 |
| *Tulasnella rosea* | MN947567 | *Spiculaea ciliata* | CLM1767 | Arifin et al. 2021 |
| *Tulasnella rosea* | MN947564 | *Spiculaea ciliata* | CLM1790 | Arifin et al. 2021 |
| *Tulasnella rosea* | MN947560 | *Spiculaea ciliata* | CLM1797 | Arifin et al. 2021 |
| *Tulasnella rosea* | MN947559 | *Spiculaea ciliata* | CLM1798 | Arifin et al. 2021 |
| *Tulasnella rosea* | MN947558 | *Spiculaea ciliata* | CLM1820 | Arifin et al. 2021 |
| *Tulasnella rosea* | MN947557 | *Spiculaea ciliata* | CLM1847 | Arifin et al. 2021 |
| *Tulasnella rosea* | MN947556 | *Spiculaea ciliata* | CLM1848 | Arifin et al. 2021 |
| *Tulasnella rosea* | MN947555 | *Spiculaea ciliata* | CLM1802 | Arifin et al. 2021 |
| *Tulasnella rosea* | MN947554 | *Spiculaea ciliata* | CLM1804 | Arifin et al. 2021 |
| *Tulasnella rosea* | MN947553 | *Spiculaea ciliata* | CLM1814 | Arifin et al. 2021 |
| *Tulasnella rosea* | MN947552 | *Spiculaea ciliata* | CLM1827 | Arifin et al. 2021 |
| *Tulasnella rosea* | MN947551 | *Spiculaea ciliata* | CLM1828 | Arifin et al. 2021 |
| *Tulasnella rosea* | MN947550 | *Spiculaea ciliata* | CLM1829 | Arifin et al. 2021 |
| *Tulasnella rosea* | MN947549 | *Spiculaea ciliata* | CLM1830 | Arifin et al. 2021 |

| | | | | |
|---|---|---|---|---|
| *Tulasnella rosea* | MN947548 | *Spiculaea ciliata* | CLM1831 | Arifin et al. 2021 |
| *Tulasnella rosea* | MN947547 | *Spiculaea ciliata* | CLM1844 | Arifin et al. 2021 |
| *Tulasnella* sp. ECU6 | KC152401 | Fallen branch | DC185 | Cruz et al. 2014 |
| *Tulasnella* sp. ECU6 | KC152402 | Fallen branch | DC262 | Cruz et al. 2014 |
| *Tulasnella* sp. ECU5 | KC152397 | Fallen branch | DC225 | Cruz et al. 2014 |
| *Tulasnella* sp. ECU5 | KC152398 | Fallen branch | DC225 | Cruz et al. 2014 |
| *Tulasnella aurantiaca* | MK626568 | Rotten wood | DAOMC 252086 | Mack et al. 2021 |
| *Tulasnella aurantiaca* | MK626686 | Rotten wood | DAOMC 2521988 | Mack et al. 2021 |
| *Tulasnella aurantiaca* | MK626687 | Rotten wood | DAOMC 252083 | Mack et al. 2021 |
| *Tulasnella tomaculum* | AY373296 | Unknown | KC429 | McCormick et al. 2004 |
| *Tulasnella tomaculum* | KC152380 | Unknown | K(M)123675 | Cruz et al. 2014 |
| *Tulasnella warcupii* | MT231823 | *Arthrochilus irritabilis* | CL033.4 clone1 | This study |
| *Tulasnella warcupii* | MT231824 | *Arthrochilus irritabilis* | CL033.4 clone2 | This study |
| *Tulasnella warcupii* | MT231825 | *Arthrochilus irritabilis* | CL033.4 clone3 | This study |
| *Tulasnella warcupii* | MT231826 | *Arthrochilus irritabilis* | CL033.4 clone4 | This study |
| *Tulasnella warcupii* | MT231827 | *Arthrochilus irritabilis* | CL033.4 clone5 | This study |
| *Tulasnella warcupii* | MT214497 | *Arthrochilus irritabilis* | CLM1674 | This study |
| *Tulasnella warcupii* | MT214498 | *Arthrochilus irritabilis* | CLM1694 | This study |
| *Tulasnella warcupii* | MT214499 | *Arthrochilus irritabilis* | CLM1696 | This study |
| *Tulasnella warcupii* | MT214500 | *Arthrochilus irritabilis* | CLM1700 | This study |
| *Tulasnella warcupii* | MT214501 | *Arthrochilus irritabilis* | CLM1701 | This study |
| *Tulasnella warcupii* | N/A | *Arthrochilus irritabilis* | AI002-6 | This study |
| *Tulasnella warcupii* | MT231828 | *Arthrochilus prolixus* | CL152.1 clone1 | This study |
| *Tulasnella warcupii* | MT231831 | *Arthrochilus prolixus* | CL152.1 clone3 | This study |
| *Tulasnella warcupii* | MT231830 | *Arthrochilus prolixus* | CL152.1 clone5 | This study |
| *Tulasnella warcupii* | MT231829 | *Arthrochilus prolixus* | CL152.1 clone2 | This study |
| *Tulasnella warcupii* | MT231832 | *Arthrochilus prolixus* | CL152.1 clone4 | This study |
| *Tulasnella warcupii* | MT214506 | *Arthrochilus prolixus* | CLM2242 | This study |
| *Tulasnella warcupii* | MT214502 | *Arthrochilus prolixus* | CLM2243 | This study |
| *Tulasnella warcupii* | MT214540 | *Arthrochilus latipes* | CLM2236 | This study |
| *Tulasnella warcupii* | MT214541 | *Arthrochilus latipes* | CLM2238 | This study |
| *Tulasnella warcupii* | MT231871 | *Arthrochilus latipes* | CL130.3 clone1 | This study |
| *Tulasnella warcupii* | MT231872 | *Arthrochilus latipes* | CL130.3 clone2 | This study |
| *Tulasnella warcupii* | MT231833 | *Arthrochilus latipes* | CL130.2 clone1 | This study |
| *Tulasnella warcupii* | MT231834 | *Arthrochilus latipes* | CL130.2 clone2 | This study |
| *Tulasnella warcupii* | MT231835 | *Arthrochilus latipes* | CL130.2 clone3 | This study |
| *Tulasnella warcupii* | MT231836 | *Arthrochilus latipes* | CL130.2 clone4 | This study |
| *Tulasnella warcupii* | MT214503 | *Arthrochilus latipes* | CLM2239 | This study |
| *Tulasnella warcupii* | MT214504 | *Arthrochilus latipes* | CLM2240 | This study |
| *Tulasnella warcupii* | MT214505 | *Arthrochilus latipes* | CLM2237 | This study |
| *Tulasnella warcupii* | MT231840 | *Paracaleana nigrita* | CL103.2 clone1 | This study |
| *Tulasnella warcupii* | MT231837 | *Paracaleana nigrita* | CL103.2 clone2 | This study |

| | | | | |
|---|---|---|---|---|
| *Tulasnella warcupii* | MT231838 | *Paracaleana nigrita* | CL103.2 clone3 | This study |
| *Tulasnella warcupii* | MT231839 | *Paracaleana nigrita* | CL103.4 clone4 | This study |
| *Tulasnella warcupii* | MT231841 | *Arthrochilus rosulatus* | CL154.1 clone1 | This study |
| *Tulasnella warcupii* | MT231842 | *Arthrochilus rosulatus* | CL154.1 clone2 | This study |
| *Tulasnella warcupii* | MT231843 | *Arthrochilus rosulatus* | CL154.1 clone3 | This study |
| *Tulasnella warcupii* | MT231844 | *Arthrochilus rosulatus* | CL154.2 clone1 | This study |
| *Tulasnella warcupii* | MT231845 | *Arthrochilus rosulatus* | CL154.3 clone1 | This study |
| *Tulasnella warcupii* | MT214528 | *Arthrochilus rosulatus* | CLM2249 | This study |
| *Tulasnella warcupii* | MT214507 | *Arthrochilus rosulatus* | CLM2250 | This study |
| *Tulasnella warcupii* | MT214527 | *Arthrochilus rosulatus* | CLM2251 | This study |
| *Tulasnella warcupii* | MT214508 | *Arthrochilus rosulatus* | CLM2252 | This study |
| *Tulasnella warcupii* | MT214509 | *Arthrochilus rosulatus* | CLM2253 | This study |
| *Tulasnella warcupii* | MT214529 | *Arthrochilus rosulatus* | CLM2254 | This study |
| *Tulasnella warcupii* | MT214510 | *Arthrochilus rosulatus* | CLM2255 | This study |
| *Tulasnella warcupii* | MT214530 | *Arthrochilus rosulatus* | CLM2257 | This study |
| *Tulasnella warcupii* | MT214511 | *Arthrochilus rosulatus* | CLM2258 | This study |
| *Tulasnella warcupii* | MT231846 | *Paracaleana disjuncta* | CL075.2 clone2 | This study |
| *Tulasnella warcupii* | MT231866 | *Paracaleana disjuncta* | CL075.2 clone3 | This study |
| *Tulasnella warcupii* | MT214512 | *Arthrochilus dockrillii* | CLM2259 | This study |
| *Tulasnella warcupii* | MT214513 | *Arthrochilus dockrillii* | CLM2263 | This study |
| *Tulasnella warcupii* | MT231847 | *Arthrochilus dockrillii* | CL155.3 clone1 | This study |
| *Tulasnella warcupii* | MT231848 | *Arthrochilus dockrillii* | CL155.4 clone2 | This study |
| *Tulasnella warcupii* | MT214514 | *Arthrochilus dockrillii* | CLM2260 | This study |
| *Tulasnella warcupii* | MT214515 | *Arthrochilus dockrillii* | CLM2261 | This study |
| *Tulasnella warcupii* | MT214516 | *Arthrochilus dockrillii* | CLM2262 | This study |
| *Tulasnella warcupii* | MT231849 | *Arthrochilus dockrillii* | CL155.2 clone1 | This study |
| *Tulasnella warcupii* | MT231852 | *Arthrochilus dockrillii* | CL155.2 clone2 | This study |
| *Tulasnella warcupii* | MT231853 | *Arthrochilus dockrillii* | CL155.2 clone3 | This study |
| *Tulasnella warcupii* | MT231850 | *Arthrochilus dockrillii* | CL155.4 clone1 | This study |
| *Tulasnella warcupii* | MT231851 | *Arthrochilus dockrillii* | CL155.4 clone3 | This study |
| *Tulasnella warcupii* | MT231854 | *Arthrochilus dockrillii* | CL156.1 clone3 | This study |
| *Tulasnella warcupii* | MT231855 | *Arthrochilus dockrillii* | CL156.2 clone1 | This study |
| *Tulasnella warcupii* | MT231856 | *Arthrochilus dockrillii* | CL156.2 clone2 | This study |
| *Tulasnella warcupii* | MT231857 | *Arthrochilus dockrillii* | CL156.3 clone1 | This study |
| *Tulasnella warcupii* | MT231858 | *Arthrochilus dockrillii* | CL156.3 clone2 | This study |
| *Tulasnella warcupii* | MT231859 | *Arthrochilus dockrillii* | CL156.1 clone1 | This study |
| *Tulasnella warcupii* | MT231869 | *Arthrochilus dockrillii* | CL156.1 clone2 | This study |
| *Tulasnella warcupii* | MT231870 | *Arthrochilus dockrillii* | CL156.2 clone3 | This study |
| *Tulasnella warcupii* | MT214538 | *Arthrochilus dockrillii* | CLM2264 | This study |
| *Tulasnella warcupii* | MT214539 | *Arthrochilus dockrillii* | CLM2265 | This study |
| *Tulasnella warcupii* | MT214517 | *Arthrochilus dockrillii* | CLM2266 | This study |
| *Tulasnella warcupii* | KF476596 | *Arthrochilus oreophilus* | CLM027 | Linde et al. 2014 |
| *Tulasnella warcupii* | KF476601 | *Arthrochilus oreophilus* | CLM022 | Linde et al. 2014 |

| | | | | |
|---|---|---|---|---|
| *Tulasnella warcupii* | KF476600 | *Arthrochilus oreophilus* | CLM007 | Linde et al. 2014 |
| *Tulasnella warcupii* | KF476598 | *Arthrochilus oreophilus* | CLM091 | Linde et al. 2014 |
| *Tulasnella warcupii* | KF476597 | *Arthrochilus oreophilus* | CLM092 | Linde et al. 2014 |
| *Tulasnella warcupii* | KF476599 | *Arthrochilus oreophilus* | CLM028 | Linde et al. 2014 |
| *Tulasnella warcupii* | MT214532 | *Arthrochilus oreophilus* | CLM2230 | This study |
| *Tulasnella warcupii* | MT214533 | *Arthrochilus oreophilus* | CLM2231 | This study |
| *Tulasnella warcupii* | MT214534 | *Arthrochilus oreophilus* | CLM2232 | This study |
| *Tulasnella warcupii* | MT214535 | *Arthrochilus oreophilus* | CLM2233 | This study |
| *Tulasnella warcupii* | MT214536 | *Arthrochilus oreophilus* | CLM2234 | This study |
| *Tulasnella warcupii* | MT214537 | *Arthrochilus oreophilus* | CLM2235 | This study |
| *Tulasnella warcupii* | MT214531 | *Arthrochilus oreophilus* | CLM2277 | This study |
| *Tulasnella warcupii* | MT214523 | *Arthrochilus oreophilus* | CLM2268 | This study |
| *Tulasnella warcupii* | MT214524 | *Arthrochilus oreophilus* | CLM2275 | This study |
| *Tulasnella warcupii* | MT214526 | *Arthrochilus oreophilus* | CLM2276 | This study |
| *Tulasnella warcupii* | MT214525 | *Arthrochilus oreophilus* | CLM2229 | This study |
| *Tulasnella warcupii* | MT231864 | *Arthrochilus oreophilus* | CL158.2 clone1 | This study |
| *Tulasnella warcupii* | MT231865 | *Arthrochilus oreophilus* | CL158.3 clone1 | This study |
| *Tulasnella warcupii* | MT231861 | *Arthrochilus oreophilus* | CL158.1 clone1 | This study |
| *Tulasnella warcupii* | MT231862 | *Arthrochilus oreophilus* | CL158.1 clone2 | This study |
| *Tulasnella warcupii* | MT231863 | *Arthrochilus oreophilus* | CL158.2 clone2 | This study |
| *Tulasnella warcupii* | MT214518 | *Arthrochilus stenophyllus* | CLM2269 | This study |
| *Tulasnella warcupii* | MT214519 | *Arthrochilus stenophyllus* | CLM2271 | This study |
| *Tulasnella warcupii* | MT214520 | *Arthrochilus stenophyllus* | CLM2272 | This study |
| *Tulasnella warcupii* | MT214521 | *Arthrochilus stenophyllus* | CLM2273 | This study |
| *Tulasnella warcupii* | MT214522 | *Arthrochilus stenophyllus* | CLM2274 | This study |
| *Tulasnella warcupii* | MT231867 | *Arthrochilus stenophyllus* | CL157.2 clone1 | This study |
| *Tulasnella warcupii* | MT231860 | *Arthrochilus stenophyllus* | CL157.2 clone2 | This study |
| *Tulasnella warcupii* | MT231868 | *Arthrochilus stenophyllus* | CL157.2 clone3 | This study |
| OTU 3 | KF476594 | *Arthrochilus oreophilus* | CLM084 | Linde et al. 2014 |
| OTU 3 | KF476595 | *Arthrochilus oreophilus* | CLM085 | Linde et al. 2014 |
| *Tulasnella australiensis* | KF476602 | *Arthrochilus oreophilus* | CLM031 | Linde et al. 2014 |
| *Tulasnella australiensis* | OM311924 | *Arthrochilus rosulatus* | CL154.2 clone3 | This study |
| *Tulasnella australiensis* | MT003751 | *Cryptostylis leptochila* | CLM1670 | Arifin et al. 2021 |
| *Tulasnella australiensis* | MT003731 | *Cryptostylis erecta* | CLM1944 | Arifin et al. 2021 |
| *Tulasnella australiensis* | MT003730 | *Cryptostylis erecta* | CLM1945 | Arifin et al. 2021 |
| *Tulasnella australiensis* | MT003729 | *Cryptostylis subulata* | CLM1946 | Arifin et al. 2021 |
| *Tulasnella australiensis* | MT003728 | *Cryptostylis subulata* | CLM1947 | Arifin et al. 2021 |
| *Tulasnella australiensis* | MT003747 | *Cryptostylis leptochila* | CLM2073 | Arifin et al. 2021 |
| OTU 2 | OM311925 | *Arthrochilus oreophilus* | CL112.2 clone1 | This study |
| *Tulasnella secunda* | KF476586 | *Drakaea glyptodon* | CLM258 | Phillips et al. 2011 |
| *Tulasnella secunda* | KF476583 | *Drakaea glyptodon* | CLM257 | Phillips et al. 2011 |
| *Tulasnella secunda* | KF476577 | *Drakaea glyptodon* | CLM259 | Phillips et al. 2011 |

| | | | | |
|---|---|---|---|---|
| *Tulasnella secunda* | HQ386743 | *Drakaea glyptodon* | RP-2011 | Phillips et al. 2011 |
| *Tulasnella secunda* | KF476590 | *Drakaea livida* | CLM255 | Phillips et al. 2011 |
| *Tulasnella secunda* | KF476576 | *Drakaea livida* | CLM273 | Phillips et al. 2011 |
| *Tulasnella secunda* | HQ386778 | *Drakaea livida* | RP-2011 | Phillips et al. 2011 |
| *Tulasnella secunda* | HQ386747 | *Drakaea livida* | RP-2011 | Phillips et al. 2011 |
| *Tulasnella secunda* | KF476579 | *Drakaea confluens* | CLM266 | Phillips et al. 2011 |
| *Tulasnella secunda* | KF476592 | *Drakaea confluens* | CLM253 | Phillips et al. 2011 |
| *Tulasnella secunda* | KF476591 | *Drakaea gracilis* | CLM261 | Phillips et al. 2011 |
| *Tulasnella secunda* | KF476578 | *Drakaea gracilis* | CLM277 | Phillips et al. 2011 |
| *Tulasnella secunda* | KF476575 | *Drakaea elastica* | CLM009 | Phillips et al. 2011 |
| *Tulasnella secunda* | KF476593 | *Drakaea elastica* | CLM260 | Phillips et al. 2011 |
| *Tulasnella secunda* | HQ386750 | *Drakaea elastica* | RP-2011 | Phillips et al. 2011 |
| *Tulasnella secunda* | KF476570 | *Drakaea elastica* | CLM228 | Phillips et al. 2011 |
| *Tulasnella secunda* | KF476585 | *Drakaea isolata* | CLM276 | Phillips et al. 2011 |
| *Tulasnella secunda* | KF476589 | *Drakaea concolor* | CLM252 | Phillips et al. 2011 |
| *Tulasnella secunda* | KF476588 | *Drakaea concolor* | CLM251 | Phillips et al. 2011 |
| *Tulasnella secunda* | HQ386752 | *Drakaea micrantha* | RP-2011 | Phillips et al. 2011 |
| *Tulasnella secunda* | HQ386753 | *Drakaea micrantha* | RP-2011 | Phillips et al. 2011 |
| *Tulasnella secunda* | HQ386754 | *Drakaea micrantha* | RP-2011 | Phillips et al. 2011 |
| *Tulasnella secunda* | HQ386763 | *Drakaea micrantha* | RP-2011 | Phillips et al. 2011 |
| *Tulasnella secunda* | HQ386756 | *Drakaea thynniphila* | RP-2011 | Phillips et al. 2011 |
| *Tulasnella secunda* | HQ386761 | *Drakaea thynniphila* | RP-2011 | Phillips et al. 2011 |
| *Tulasnella secunda* | HQ386762 | *Drakaea thynniphila* | RP-2011 | Phillips et al. 2011 |
| *Tulasnella secunda* | KF476584 | *Paracaleana hortiorum* | CLM267 | Linde et al. 2014 |
| *Tulasnella secunda* | OM311926 | *Paracaleana hortiorum* | CL171.1 clone1 | This study |
| *Tulasnella secunda* | OM311927 | *Paracaleana hortiorum* | CL171.2 clone1 | This study |
| *Tulasnella secunda* | OM311930 | *Caleana major* | CL108 clone2 | This study |
| *Tulasnella secunda* | OM311931 | *Caleana major* | CL108 clone3 | This study |
| *Tulasnella secunda* | KF476574 | *Paracaleana terminalis* | CLM268 | Linde et al. 2014 |
| *Tulasnella secunda* | ON256292 | *Paracaleana terminalis* | CLM2212 | This study |
| *Tulasnella secunda* | ON256293 | *Paracaleana terminalis* | CLM2214 | This study |
| *Tulasnella secunda* | ON256295 | *Paracaleana terminalis* | CLM2217 | This study |
| *Tulasnella secunda* | ON256296 | *Paracaleana terminalis* | CLM2218 | This study |
| *Tulasnella secunda* | ON256297 | *Paracaleana terminalis* | CLM2219 | This study |
| *Tulasnella secunda* | ON256294 | *Paracaleana terminalis* | CLM2215 | This study |
| *Tulasnella secunda* | ON256299 | *Paracaleana terminalis* | CLM2216 | This study |
| *Tulasnella secunda* | ON256298 | *Paracaleana terminalis* | CLM2213 | This study |
| *Tulasnella secunda* | KF476580 | *paracaleana triens* | CLM274 | Linde et al. 2014 |
| *Tulasnella secunda* | KF476568 | *Paracaleana minor* | CLM222 | Linde et al. 2014 |
| *Tulasnella secunda* | OM311928 | *Paracaleana disjuncta* | CL075.2 clone1 | This study |
| *Tulasnella secunda* | OM311929 | *Paracaleana disjuncta* | CL075.2 clone4 | This study |
| *Tulasnella secunda* | KF476573 | *Paracaleana lyonsii* | CLM272 | Linde et al. 2014 |
| *Tulasnella secunda* | ON256301 | *Paracaleana nigrita* | CLM2224 | This study |

| | | | | |
|---|---|---|---|---|
| *Tulasnella secunda* | ON256302 | *Paracaleana nigrita* | CLM2225 | This study |
| *Tulasnella secunda* | ON256305 | *Paracaleana nigrita* | CLM2226 | This study |
| *Tulasnella secunda* | ON256303 | *Paracaleana nigrita* | CLM2228 | This study |
| *Tulasnella nerrigaensis* | MW575584 | *Calochilus robertsonii* | CLM1869 | Arifin et al. 2022 |
| *Tulasnella nerrigaensis* | MW575585 | *Calochilus robertsonii* | CLM1870 | Arifin et al. 2022 |
| *Tulasnella nerrigaensis* | MW575589 | *Calochilus robertsonii* | CLM1875 | Arifin et al. 2022 |
| *Tulasnella nerrigaensis* | MW575587 | *Calochilus robertsonii* | CLM1873 | Arifin et al. 2022 |
| *Tulasnella nerrigaensis* | MW575588 | *Calochilus robertsonii* | CLM1874 | Arifin et al. 2022 |
| *Tulasnella violea* | KC152437 | Decayed wood | DC293 | Cruz et al. 2014 |
| *Tulasnella violea* | KC152415 | Fallen branch | DC177 | Cruz et al. 2014 |
| OTU 4 | MZ576537 | *Arthrochilus huntianus* | CL045 clone1 | This study |
| OTU 4 | MZ576538 | *Arthrochilus huntianus* | CL045 clone2 | This study |
| OTU 4 | MZ576539 | *Arthrochilus huntianus* | CL045 clone3 | This study |
| OTU 4 | MZ576540 | *Arthrochilus huntianus* | CL045 clone8 | This study |
| OTU 4 | MZ576541 | *Arthrochilus huntianus* | CL045 clone7 | This study |
| OTU 4 | MZ576542 | *Arthrochilus huntianus* | CL045 clone4 | This study |
| OTU 4 | MZ576543 | *Arthrochilus huntianus* | CL045 clone5 | This study |
| OTU 4 | MZ576544 | *Arthrochilus huntianus* | CL045 clone6 | This study |
